# Tree Shape-based approaches for the Comparative study of Cophylogeny

**DOI:** 10.1101/388116

**Authors:** Mariano Avino, Garway T. Ng, Yiying He, Mathias S. Renaud, Bradley R. Jones, Art F. Y. Poon

**Author notes:** Correspondence to be sent to: Department of Pathology and Laboratory Medicine, Dental Sciences Building, Western University, London, ON N6A 5C1, Canada.

## Abstract

Cophylogeny is the congruence of phylogenetic relationships between two different groups of organisms due to their long-term interaction, such as between host and pathogen species. Discordance between host and pathogen phylogenies may occur due to pathogen host-switch events, pathogen speciation within a host species, and extinction. Here, we investigated the use of tree shape distance measures to quantify the degree of cophylogeny for the comparative analysis of host-pathogen interactions across taxonomic groups.

We firstly implemented a coalescent model to simulate pathogen phylogenies within a fixed host tree, given the cospeciation probability, migration rate between hosts, and pathogen speciation rate within hosts.

Next, we used simulations from this model to evaluate 13 distance metrics between these trees and the host tree, including Robinson-Foulds distance and two kernel distances that we developed for labeled and unlabeled trees, which use branch lengths and can accommodate trees of different sizes. Finally, we used these distance metrics to revisit actual datasets from published cophylogenetic studies across all taxonomic groups, where authors described the observed associations as representing a high or low degree of cophylogeny.

Our simulation analyses demonstrated that some metrics are more informative than others with respect to specific coevolution parameters. For example, the Sim metric was the most responsive to variation in coalescence rates, whereas the unlabeled kernel metric was the most responsive to cospeciation probabilities. We also determined that distance metrics were more informative about the model parameters when the underlying parameter values did not assume extreme values, *e.g.,* rapid host switching. When applied to real datasets, projection of these trees’ associations into a parameter space defined by the 13 distance metrics revealed some clustering of studies reporting low concordance. This suggested that different investigators are describing concordance in a consistent way across biological systems, and that these expert subjective assessments can be at least partly quantified using distance metrics.

Our results support the hypothesis that tree distance measures can be useful for quantifying host and pathogen cophylogeny. This motivates the usage of distance metrics in the field of coevolution and supports the development of simulation-based methods, *i.e.,* approximate Bayesian computation, to estimate coevolutionary parameters from the discordant shapes of host and pathogen trees. [tree shape; cophylogeny; codivergence; coevolution; host switching; tree metrics; kernel]

---

Coevolution occurs when two or more species exert a reciprocal influence on one another’s evolutionary trajectories (Vermeij 1994). These effects may be mediated by beneficial (mutualistic) or deleterious associations *(e.g.,* parasitism, predation). For simplicity, we will only refer to ‘host’ and ‘pathogen’ species, although we recognize that many other roles in coevolutionary interactions exist in nature. A cophylogenetic study is a comparative analysis of the evolutionary relationships within sets of host and pathogen species, and the extent that these relationships are correlated back in time. Host-pathogen associations are frequently visualized by a ‘tanglegram’, in which the associations are mapped to the two phylogenies by drawing association edges between the respective host and pathogen taxa (Page 1993). If the topologies of the two phylogenies are fully concordant, then there exists an arrangement of their branches (by rotation around ancestral nodes) such that the association edges do not intersect — the trees are completely ‘untangled’. This situation implies that the interactions between the host and pathogen species are so strong that the diversification of the pathogen species is entirely constrained by that of their hosts. Discordant trees can also yield an untangled graph. Thus, the number of intersecting association edges is a more useful metric for optimizing visual layouts than for inferring biological processes.

Any single tanglegram may be explained by a large number of different combinations of events in the past, including pathogen extinction, host switching, incomplete lineage sorting (Pamilo and Nei 1988), pathogen speciation / duplication, and unobserved species; see Charleston and Perkins (2006) for a detailed discussion of these event types. Increasing numbers of events in the coevolutionary history of the host and pathogen species will tend to result in a lower degree of topological concordance between their phylogenies. Estimating the optimal reconstruction of such events to explain the present-day associations between the tip taxa of the host and pathogen phylogenies is known as reconciliation inference (Doyon et al. 2011). A well-characterized approach to reconciliation inference is to assign a cost to each type of event and to identify the most parsimonious (minimum cost) distribution of events. However, the resulting solution is sensitive to the investigator’s choice of costs and becomes exceedingly difficult for larger trees — indeed, this approach has formally been determined to be a computationally intractable (NP-complete) problem (Ovadia et al. 2011). On the other hand, it is not uncommon to display the tanglegram and make a qualitative, subjective assessment about the extent of cospeciation. We propose to develop methods that occupy the middle-ground between these extremes, using simple quantitative methods to estimate coevolutionary parameters.

There is an abundance of quantitative methods for comparing trees with respect to their topology and/or branch lengths. For example, numerous investigators have proposed various summary statistics that each extract certain characteristics of tree shapes, such as asymmetry (*e.g.*, Colless’ index) and thereby reduce the tree to a single number; for a comprehensive review, see Mooers (1997). Summary statistics provide a convenient framework for comparing trees, which are otherwise statistically complex objects. However, many of these statistics are difficult to normalize to differences in tree size (Stam 2002), and can be strongly influenced by sampling for rapidly-evolving taxa (Dearlove and Frost 2015). In addition the inherent dimensionality reduction of these summary statistics is often accompanied by a critical loss of information about the underlying biological processes, which can limit the utility of any one statistic. For this reason, recent studies have begun to use feature selection methods to find optimal combinations of summary statistics (*e.g.*, Saulnier et al. 2017).

Where a summary statistic maps a tree to a number, a distance metric maps two trees to a number that quantifies their level of discordance. One of the earliest distances for trees was the cophenetic correlation (Sokal and Rohlf 1962), in which the ancestral node heights for each pair of tips in the tree is represented by a distance matrix. The ordinary product-moment correlation for two trees is then calculated from the element-wise comparison of their respective matrices. A similar distance described by Williams and Clifford (1971) restricts this comparison to the internal nodes of the trees, and measures path lengths between tips by the number of nodes — following Kuhner and Yamato (2014) we refer to this distance as ‘Node’. Other distances require the tip labels on the trees to match. For instance, the Maximum Agreement Subtree (MAST; Gordon 1980) distance is based on the largest labeled subtree that is common to both trees. The Robinson-Foulds distance (RF; Robinson and Foulds 1981) counts the minimum number of operations required to transform one tree topology to another, given that they relate the same set of taxa. It is by far the most popular and cited distance for comparing trees (Table 1). Subsequently, the RF distance has been extended to incorporate branch lengths (RFL and KF; Robinson and Foulds 1979; Kuhner and Felsenstein 1994). In addition, a very similar distance metric to RF that can accommodate unrooted trees and branch lengths was proposed by Penny et al. (1982). A normalized version of this metric was later introduced by Geoghegan et al. (nPH85; 2017). Critchlow et al. (1996) described a distance (Trip) based on the number of shared subtrees relating triples of taxa, which was extended by (Kuhner and Yamato 2014) to utilize branch lengths (TripL), who also described a second metric (Int) that sums the differences in internode distances in the trees. More recently, Hein et al. (2004) proposed a distance (Sim) based on the probability that a random point in one tree is on a branch leading to the same set of tip labels in a second tree; and a distance (Align) described by Nye et al. (2005) counts the number of mismatches in the best one-to-one mapping of branches for two trees.

**Table 1:** Summary of tree distance metrics examined in this study. In addition to the kernel metrics kU and kL, we evaluated two additional metrics where branch lengths were normalized by the mean (kUn and kLn). ‘Diff. size’ indicates which distances do not require the trees to have the same numbers of tips. ‘Diff. labels’ indicates which distances do not require the trees to relate the same taxa, *i.e.,* to have the same labels. ‘Use lengths’ indicates which distances utilize the differences in branch lengths when comparing trees. We enumerated citations in the literature by querying Google Scholar (last access date, June 31, 2017) for papers associated with the respective distance metrics and software, and then filtered the results for coevolutionary studies (‘Coevol.’).

| Distance | Reference | Diff.<br>size | Diff.<br>labels | Use<br>lengths | Citations<br>Total | Coevol. |
| --- | --- | --- | --- | --- | --- | --- |
| RF | Robinson and Foulds (1981) |  |  |  | 1561 | 3 |
| nPH85 | (Penny et al. 1982; Geoghegan et al. 2017) |  |  | Y | 227 | 1 |
| Trip | Critchlow et al. (1996) | Y |  |  | 103 | 0 |
| MAST | Gordon (1980) |  | Y |  | 69 | 0 |
| Align | Nye et al. (2005) |  |  |  | 123 | 1 |
| Node | Williams and Clifford (1971) |  | Y |  | 82 | 0 |
| RFL/KF | Robinson and Foulds (1979);<br>Kuhner and Felsenstein (1994) |  |  | Y | 725 | 0 |
| Sim | Hein et al. (2004) | Y |  | Y | 540 | 0 |
| TripL | Kuhner and Yamato (2014) |  |  | Y | 13 | 0 |
| kU | Poon et al. (2013) | Y | Y | Y | 22 | 0 |
| kL | This study | Y |  | Y | n/a |  |

The majority of these distance metrics can be computed efficiently, and several utilize branch lengths in addition to tree topologies (Table 1). On the other hand, most of the distances require the trees to have the same numbers of tips and the same tip labels, **e.g.*,* relating the same taxa. In previous work, we proposed a new tree distance measure (Poon et al. 2013) based on a kernel function from computational linguistics (Moschitti 2006) that essentially counts the number of isomorphic fragments shared by two trees, while penalizing fragments for their discordance in branch lengths. The resulting distance measure is normalized for differences in tree sizes and can optionally ignore tip labels, such that it can be applied to trees relating different sets of taxa.

A distance metric may be difficult to interpret without some absolute scale or reference distribution. Thus, the discordance between host and pathogen trees can also be quantified by an independence test (De Vienne et al. 2013), which evaluates the probability that an equal or shorter distance is obtained by chance given a null distribution. Hence, this test essentially maps the distance metric to a more interpretable scale. The null distribution can be either generated at random from the simulation of trees given a parametric model, or by the non-parametric permutation of the host and pathogen trees. Finally, we note that this is not a comprehensive review of distance metrics on trees; we acknowledge more recent and ongoing advances in this area in the Discussion section.

Although such distance measures have been widely utilized in the comparison of trees in both evolutionary and broader contexts, there are surprisingly few references to these measures in the cophylogeny literature (Table 1). We propose that tree distance metrics may provide a useful alternative to the visual assessment of tanglegrams or reconciliation methods that require the investigator to make one or more subjective decisions. In this study, our objective is to assess how much information these distance measures can extract about coevolutionary events from the discordance of host and pathogen phylogenies. This however requires evaluating these distances on sets of trees where the underlying cophylogeny process is known with absolute certainty that reconciliation methods cannot provide. Thus, we developed a reverse-time simulation framework for generating pathogen trees along a fixed host tree, from the tips towards the root, for given rates of cospeciation, duplication and host switching. This work provides a critical quantitative assessment on the potential utility of distance metrics for cophylogenetic studies, and provides detailed guidance for choosing among those metrics given prior information on the relative importance of different coevolutionary events, or to focus on specific metrics are more informative than others about coevolutionary processes.

## Methods

### Simulation methods

To simulate pathogen trees within host trees, we implemented a coalescent (reverse-time) simulation method with a custom Python script. The required inputs of this script are: (1) a Newick string representation of the host tree, with branches scaled in units of real time; (2) the coalescence rate of two pathogen lineages within the same host, Λ; (3) the migration rate for pathogen lineages between hosts (host switching), *M*, and; (4) the probability of cospeciation, *P*. The coalescence of host species was a non-random event determined by the input tree. Stochastic events were simulated using the standard Gillespie method (Gillespie 1977). The total rate of stochastic events was:

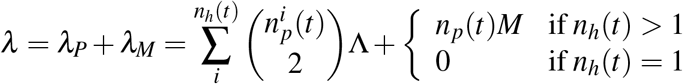

where *n_h_*(*t*) is the number of extant host lineages at time *t*, *n_p_(t*) is the number of extant pathogen lineages, and 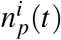 is the number of extant pathogen lineages within the i-th host. The coalescence of pathogen lineages within a host is the reverse-time analog of a duplication event, *i.e*., the speciation of a pathogen lineage into two derived species within the same host lineage. We assumed that host switching was a random process that occurred at a constant and uniform rate for any single pathogen lineage. If there was only one extant host lineage at time *t*, then we assumed that the total host switching rate *λ_M_* was effectively zero.

The simulation was initialized at the most recent tips of the host tree (*t* = 0), with a single pathogen lineage assigned to each sampled host. We did not require all host species to be sampled at the same time. The heights (relative to the most recent tip at *t* = 0) of host species as determined by the input tree are denoted as 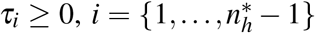, where 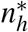 is the total number of tips in the host tree. Moreover, the sampling times of host species determined by the input tree are denoted 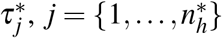, where 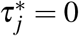 for at least one value of j. The simulation was updated iteratively back in time with a random sequence of events on the time scale of the input tree. The waiting time until the next event was drawn from an exponential distribution, Δ*t* ~ exp(λ). If the waiting time exceeded the time interval to the next highest host node *τ*, then we updated the vector of extant host nodes and reset the simulation time. If the next highest host node was a tip, then we set 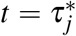 and incremented *n_p_* (*τ_j_*) by one. Otherwise if the next highest host node was an internal node, then we set *t* = *τ_i_*. All pathogen lineages carried by the affected host lineages were transferred to the ancestral host lineage, with a cospeciation probability *P* of two randomly selected lineages from the respective derived hosts being coalesced into a single ancestral lineage.

If the waiting time does not exceed the time interval to the next-highest host node *τ*, then we determined whether the next event was a host-switch or a within-host coalescence of pathogen lineages. If a host-switch event occurred with probability *λ_M_/λ*, then we selected an extant pathogen lineage at random from n_p_ and reassigned this lineage to an extant host drawn at random from *n_h_* excluding the original host. We made the simplifying assumption that all host switching events were ‘complete’, in that migration to another host species was followed by extinction in the original host. A pathogen coalescence event otherwise occurred with probability *λ_P_/λ* (= 1 – *λ_M_/λ*), in which we selected a pair of lineages occupying the same host at random to coalesce into a single ancestral lineage. Thus, the specific migration and coalescence events were uniform across pathogen lineages and pairs of lineages, respectively. Subsequently, we incremented the simulation time t by the waiting time Δt and drew the next waiting time. The simulation halts when the number of extant hosts returns to one and all tips in the host tree have been sampled. If there are multiple pathogen lineages within this ancestral host, then the simulation proceeds back in time with coalescence at a constant rate per pair until only one pathogen lineage remains.

We used the Python library *ete3* (Huerta-Cepas et al. 2016) to parse and construct tree objects. For each simulation we generated two different pathogen tree outputs: a tree in which branches were partitioned by nodes of degree-size two to record all within-host coalescence and host-switch events *(ete3* format 1); and a second tree in which this information was removed, leaving only internal nodes with degree-size 3 and terminal nodes with degree-size 1 *(ete3* format 5). Terminal node labels were subsequently removed with R package *phytools* v0.6-00 (Revell 2012). The Python script (along with data, see below) is available at our GitHub repository http://github.com/PoonLab/cophylo and released to the public domain under the GNU General Public License (version 3).

### Simulation analysis

To initialize our simulation experiments, we selected a tree relating hosts of the Hepadnaviridae (HBV) family from a recent study of host-pathogen coevolution among RNA virus families (Geoghegan et al. 2017). The authors found that the trees corresponding to this virus family and its hosts had the highest level of concordance, based on their normalized version of the Penny et al. (1982) metric (nPH85). However, the host tree Newick file published by the authors did not include any branch lengths. Consequently we obtained the time-scaled phylogenetic tree relating Metazoa published at http://timetree.org, and pruned the tree down to the host species associated with HBV, including fish, reptiles and amphibians (treated by the authors as one single host category), birds and mammals. Since a number of the distance metrics evaluated in this study required the host and pathogen trees to share the same set of tip labels, we labeled the simulated pathogen trees by their host, and initialized simulations with only one pathogen lineage per host tip.

Using this host tree, we conducted a series of ‘edge case’ simulation experiments in which two of the parameters were fixed to extreme values, and the third parameter was varied over a broad range (Table S1). The purpose of these edge case simulations was to provide outputs that were easy to interpret for validating the simulation model, and as a preliminary assessment of how the various distance metrics responded to the model parameters. To visually inspect the simulation outputs, we plotted random samples of edge case simulations alongside the host tree with DensiTree v2.2.5 (Bouckaert 2010). Next, we used Latin hypercube sampling to randomly generate 500 points that were evenly distributed in the parameter space. Specifically, we partitioned the range Λ = [0,1] into 500 intervals such that their midpoints were evenly-spaced after a log-transformation; we applied the same scheme to the range *M* = [0,1]. Since the parameter *P* is a probability instead of a range, we partitioned the range *P* = [0,1] into 500 intervals without any transformation. Next, we generated a random permutation of intervals independently along each axis, uniformly sampled one point within each cube defined by the intersection of three intervals, and simulated 100 pathogen trees using those parameter values for a total of 50,000 simulations. Correlations and mutual information (MI) tests on the performance and collinearity of metrics were performed using the R package Entropy (Hausser and Strimmer 2009).

### Data collection

Finally, we evaluated the distance metrics on phylogenies reconstructed from actual data sets. First, we collected published data sets from the literature of host and pathogen coevolution. We queried Google Scholar (https://scholar.google.ca/) on the title and abstract fields of publications with at least one of the following search terms ‘concordance’, ‘cophylogeny’ (or ‘co-phylogeny’), ‘host’, ‘pathogen’, ‘parasite’ and ‘symbiont’. Next, we manually reviewed, filtered and sorted the resulting article records into two categories (Table 2): (1) studies where the reported degree of cospeciation / codivergence, based on authors’ assessment, was moderate to high, and; (2) studies where the degree was low, with phylogenies considered too difficult to reconcile due to extensive host switching, duplication or extinction events. For each study, we obtained the sequence data using a batch query of the Genbank accession numbers. The resulting collection of 36 trees are summarized in Table 2 and are referred herein as the ‘General’ collection.

**Table 2:**
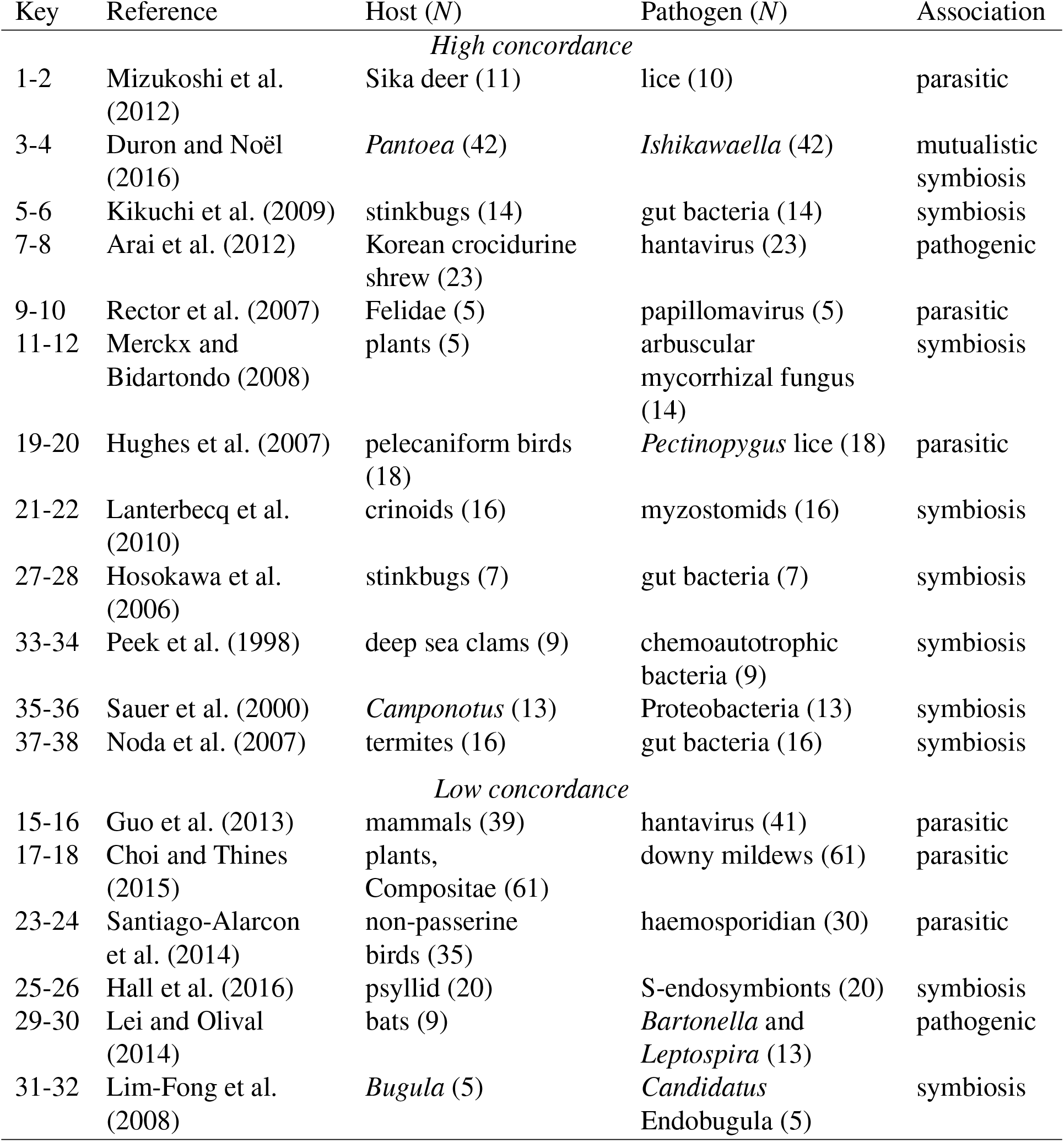
Summary of cophylogeny studies and data sets collected from the literature (the ‘General’ collection). The keys are used to map these entries to subsequent figures. *N* denotes the number of tips in the corresponding host or parasite tree.

Second, we obtained all of the 19 host-virus data sets from Geoghegan et al. (2017), which we refer to as the ‘Viral’ collection. Because the host trees in Geoghegan et al. (2017) were not available with branch lengths, we reconstructed these lengths by extracting them from time-scaled trees published at http://www.timetree.org (Hedges et al. 2006). We retrieved the Metazoa (n = 1,456 tips) and Viridiplantae (n = 373 tips) trees from this database at the taxonomic resolution of families. We mapped host species annotations from the virus phylogenies to these family-level trees using the NCBI BLAST taxonomy (Sayers et al. 2009). When more than one tip in a virus phylogeny mapped to the same host family, we collapsed those tips into a single terminal branch as in Geoghegan et al. (2017). For consistency with the original study, we applied midpoint rooting to the pathogen trees; however, we also evaluated outgroup rooting and placement of the root to minimize the distance to the host tree, but we found no significant effect on our results.

### Data processing

Because sequence alignments were not available from the studies in the ‘General’ collection, we reconstructed alignments of nucleotide or amino acid sequence data using MUSCLE (version 3.8.425, (Edgar 2004)) with the default settings. The resulting alignments were visually inspected and refined using AliView (Larsson 2014). We determined the optimal substitution model for each alignment using jModelTest 2.0 (Darriba et al. 2012) for nucleotide sequences and prottest3 (Darriba et al. 2011) for amino acid sequences, both of which employ the Akaike information criterion for model selection. Phylogenetic trees were reconstructed by maximum likelihood using PhyML 3.0 (Guindon and Gascuel 2003) and, for the ‘General’ dataset, rooted on the branches determined by the respective studies. These trees were visually inspected in FigTree (Rambaut 2009) to verify that the result was consistent with the source publications.

### Distance metrics

We used the implementation of the Robinson-Foulds (RF) distance in the R library *phangorn* v2.4.0 (Schliep 2011) with default parameters. An extension of RF (RFL/KF) incorporates branch length information into the comparison of tree topologies. We used the function KF.dist in the R library *phangorn* v2.4.0 to calculate this extended metric under the default parameters. In addition, we used the function nPH85 in R library *NELSI* v0.2 (Ho et al. 2015) to calculate the related normalized Penny-Hendy metric. To calculate the metrics Align, Node, MAST, Trip and TripL in the same framework, we ported the respective implementations from the Python script published by Kuhner and Yamato (Kuhner and Yamato 2014) into a custom R package (https://github.com/PoonLab/Kaphi).

A kernel function computes the inner product between two objects that have been mapped to a high-dimensional feature space (Aizerman et al. 1964). It is a highly efficient method for comparing complex objects for which there is a potentially enormous number of features in each object, because the kernel restricts its calculation to the comparable tiny subset of features that occur in at least one of the two objects. A larger inner product indicates that the objects share a greater number of features; hence, the kernel can be used as a measure of similarity. Poon et al. (2013) previously adapted a kernel function that operates on tree-like objects in natural language processing (Collins and Duffy 2002) to compare phylogenetic trees. The features counted by this kernel are subset trees. A subset tree is a fragment of a tree that is rooted at an ancestral node and extends down towards its descendants. It does not necessarily extend all the way to the tips of the tree – if it does, however, then it is referred to as a ‘subtree’ (Moschitti 2006). The tree shape kernel, essentially counts the number of times that subset trees with the same topology appear in both phylogenies, and then penalizes this number by the discordance in branch lengths (Poon et al. 2013). This kernel does not utilize tip labels, so we refer to it here as the unlabeled kernel distance (kU).

Furthermore, we extended the kernel method to compare subset trees on the basis of shared tip labels. We modified the recursive function used to calculate the kernel score, by substituting an indicator function **1**_*n*_1_,*n*_2__ in place of the constant 1 when the two nodes being compared are both tips (Collins and Duffy 2002; Poon et al. 2013). The function **1**_*n*_1_,*n*_2__ assumes the value 1 if *n*_1_ and *n*_2_ have the same labels, and otherwise returns 0. We refer to the resulting distance as the labeled kernel (kL). To generate kernel similarity matrices for the ‘General’ and ‘Viral’ data sets in this study, we first imported the Newick tree strings using the BioPython *phylo* module (Talevich et al. 2012). Branch lengths might subsequently be normalized by the mean branch length in each phylogeny to facilitate the comparison between host and pathogen trees with different overall rates of evolution (indicated by ‘n’ suffix for unlabeled and labeled kernels, or kUn and kLn). The kernel scores were also normalized, using the cosine method, to adjust for differences in the overall size (number of nodes) of the respective trees (Collins and Duffy 2002). Kernel principal components analysis and projections for the resulting matrices were generated using the *kernlab* package in R (Zeileis et al. 2004).

The behavior of the kernel function is controlled by several parameters. First, the branch length penalty is determined by a Gaussian radial basis function centered at zero with variance parameter σ, where a smaller σ results in a more severe penalty for subset trees with different branch lengths. Second, the kernel function includes a decay parameter λ that penalizes matching subset trees that are too large, which is useful to avoid the ‘large diagonal problem’ (Collins and Duffy 2002). Third, Moschitti (2006) introduced a parameter σ to control for subset tree matching, which we renamed s to avoid confusion with the Gaussian parameter. If *s* = 0, then matched subset trees must extend to the tips (subtrees) to be counted by the kernel. Since our trees have labels only on the tips, we fixed *s* = 0 for kL or kLn. Otherwise, the effect of labels was overwhelmed by subset tree shapes. Thus, with s = 1, the subset trees do not have to include all tips (kU or kUn). This parameter has especially significant importance for comparisons of labeled trees, because trees with congruent shapes and different sets of labels on their tips may be scored as highly similar when s = 1, and completely dissimilar when s = 0. Based on previous work (Poon et al. 2013), to evaluate the effect of these tuning parameters on the kernel function’s sensitivity and specificity for simulated data, we initiated our analyses with the default unlabeled kernel settings λ = 0.2, *σ =* 2 and s = 1. For ‘General’ and ‘Viral collection’ experiments, we also evaluated other combinations of the tuning parameters at the following values: λ = {0.1,0.3}, σ = {0.5,1,5,10,50,100}, and *s* = {0.5}.

## Results

### Edge case simulations

We implemented a backward-time (coalescent) model to simulates pathogen trees, given a fixed host tree and coevolutionary parameters: the within-host coalescence (lineage duplication) rate, Λ; migration rate, *M*; and cospeciation probability, *P*. To examine the response of different distance metrics (Table 1) to variation in these coevolutionary parameters, we initially adjusted each parameter individually while holding the others constant (Supplementary Material Table S1). The purpose of these edge case simulation experiments was to verify the expected effect of each model parameter under extreme conditions where their expected influence on pathogen tree shape was unambiguous. We also used these experiments to establish the potential for various distance measures to extract information about cospeciation processes by comparing the shapes of host and pathogen trees. To examine how pathogen tree shapes responded to changes in each model parameters, we plotted pathogen and host trees together for a small number of parameter values per edge case scenario (Fig. 1).

**Figure 1:**
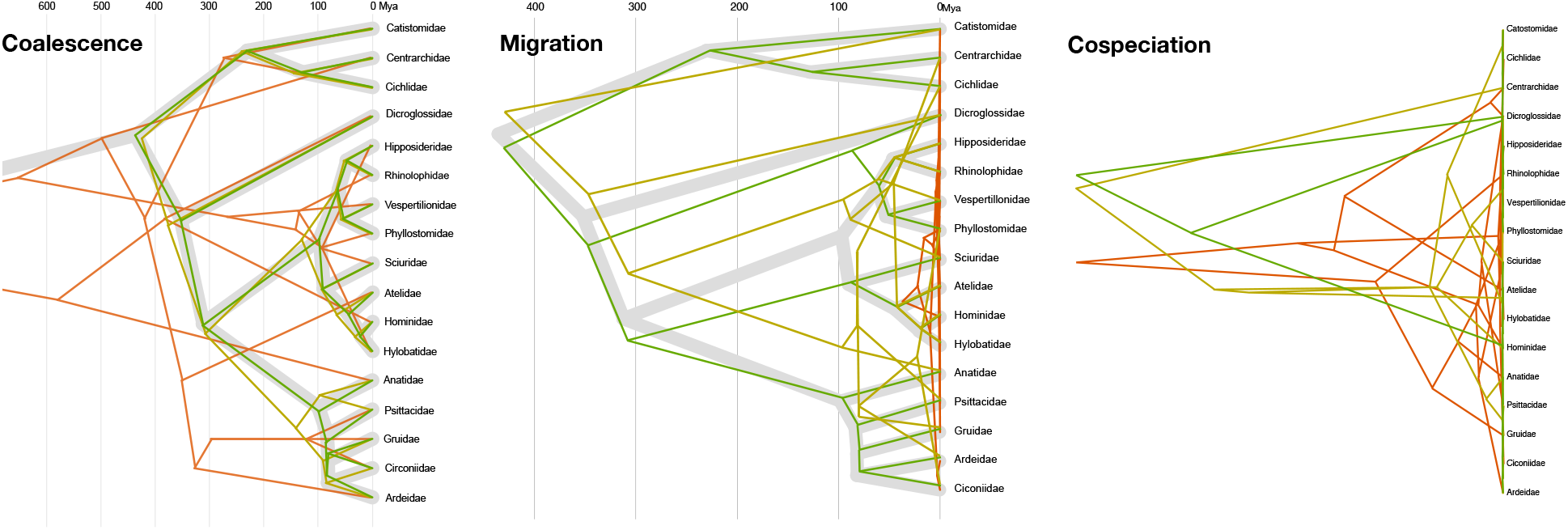
Effect of varying model parameters on simulated pathogen trees under edge cases scenarios. The host phylogeny is displayed with broad gray branches. (left) Decreasing coalescence rates Λ = {1,0.04,0.001} for green, yellow and red, respectively; *P* = 0, *M* = 0) results in a greater frequency of deep coalescent events. (centre) Increasing migration rates *M* = {0.00055,0.004375,1} for green, yellow and red, respectively; Λ = 1, *P* = 1) results in a greater frequency of host switching events. (right) Decreasing cospeciation rates *P* = {1,0.50,0.25} for green, yellow and red, respectively; Λ = 10^−6^, *M* = 0) results in a greater frequency of deep coalescent events.

Under the ‘coalescent only’ scenario, we varied the coalescence rate Λ while fixing the migration rate *M* to 10^-6^ and cospeciation probability *P* to 0. Decreasing Λ led to a greater chance of duplication events where multiple pathogen lineages co-exist in an ancestral host species (Fig. 1, left panel). Conversely, high values of Λ resulted in high concordance between pathogen and host trees. In the ‘migration only’ scenario, we varied *M* and fixed *P* to 0 and Λ to 1, the highest rate that we evaluated in this study. Increasing *M* resulted in greater discordance in shape between the host and pathogen trees as pathogen lineages switched into other hosts and immediately coalesced with the extant pathogen species, which also compressed the time-scale of the pathogen tree (Fig. 1, central panel). Finally, we varied P in the ‘cospeciation only’ scenario with *M* set to 0 and Λ set to 10^—6^. Setting Λ to the lowest value exaggerated the effect of reducing *P*, since any pathogen lineages that did not cospeciate with the host became free to coalesce on a much longer time scale (Fig. 1, right panel).

Next, we evaluated the response of the various distance metrics to individually varying the parameters within each of these three scenarios (Fig. 2). We observed substantial variation among the different tree distance measures (each scaled to their respective empirical range) in response to the within-host coalescence rate, Λ (Fig. 2, left panel), migration rate (Fig. 2, central panel) and cospeciation rate (Fig. 2, right panel). We characterized this variation by the approximate Λ, *M* and *P* values where the trends crossed a scaled distance of 0.5 (Λ_50_, *M*_50_, *P*_50_, respectively). For the majority of distance measures (RF, nPH85, MAST, Align, Node, kLn, kUn), the Λ_50_ was about 0.02/pair/Ma. The unnormalized kernel measures kU and kL were more responsive at higher coalescence rates (Λ_50_ ≈ 0.4). Trip and Sim were responsive to lower rates (Λ_50_ ≈ 0.003) and TripL and KF changed only when Λ was very low (Λ_50_ ≈ 3.5 × 10^-6^). Most of the distances displayed an approximately monotonic relationship with Λ except for Node and Align, which both increased in distance as Λ approached 1. Node, Align and KF were more responsive to slightly lower rates of migration (*M*_50_ = 10^-4^) than the other distances. All the distance metrics displayed a monotonic relationship with *M*, with the exception of TripL, Align, Node and KF, which switched around *M*=0.01/lineage/Ma. Finally, all metrics were more responsive to higher values of *P* (0.8 — 1.0) but with more variation in their response to this parameter than *M*. For example, Sim, Node and kUn sharply declined as *P* approached 1, whereas the other metrics displayed a more gradual decline with *P*. kL was the only metric that displayed a roughly linear decrease of scaled distance with *P*.

**Figure 2.**
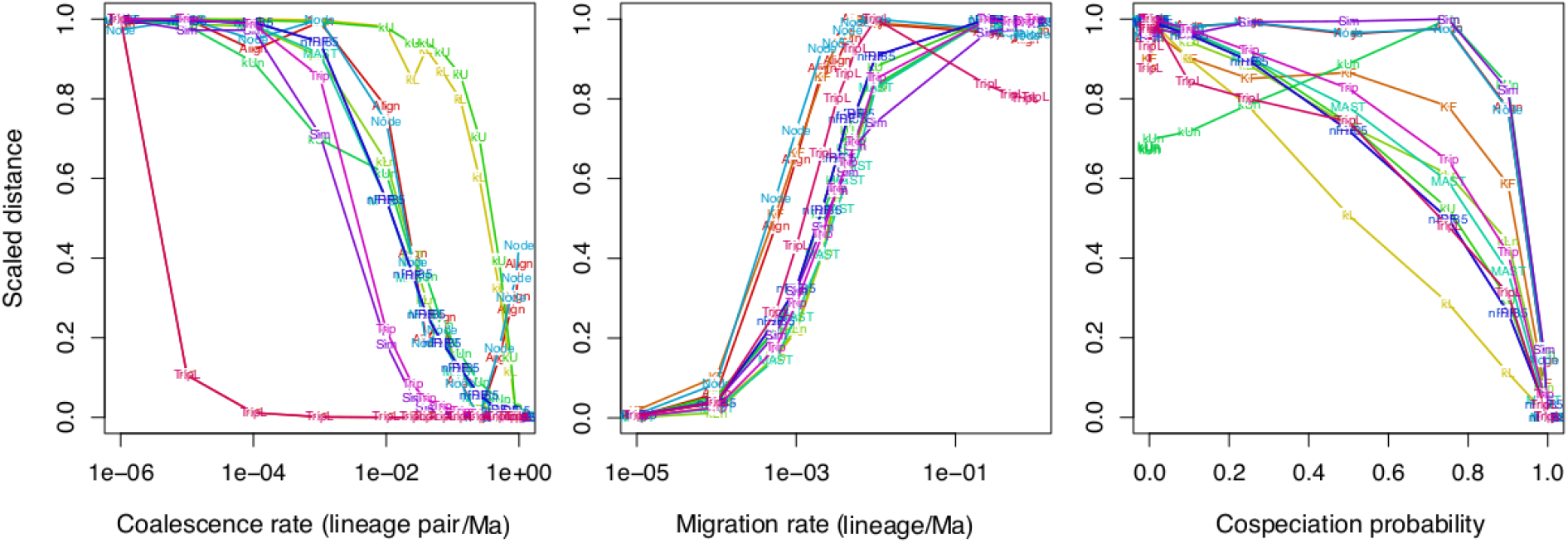
Summary of associations between tree distance measures and model parameters. Each point represents the mean distance measure (text label) for 100 replicate simulations, rescaled to range from 0 to 1 to facilitate comparisons between different measures.

### Simulation - hypercube sampling

For 500 different points sampled evenly from the parameter space defined by Λ, *M* and *P*, we used the nested coalescent model to simulate 100 pathogen trees on the phylogeny relating hosts of viruses in the HBV family, for a total of 50,000 simulations. Next, we computed the distance measures in Table 1 for every simulated tree to the ‘observed’ host tree, and averaged the distances for each of the 500 parameter settings.

Figure 3 summarizes the nonparametric (Spearman’s rank-order) correlation tests for all pairs of distance measures. We observed strong correlations (*ρ* > 0.95) among the distances kLn, RF, Trip, nPH85 and MAST. In particular, we found very high correlations between KF and TripL (*ρ* = 1), Align and Node (*ρ* = 0.94), and the two unnormalized kernels kU and kL, (*ρ* = 0.94). Interestingly, Trip and Sim, which had the same response in edge case experiments for Λ and *M*, did not have a strong positive correlation (*ρ* = 0.62).

**Figure 3:**
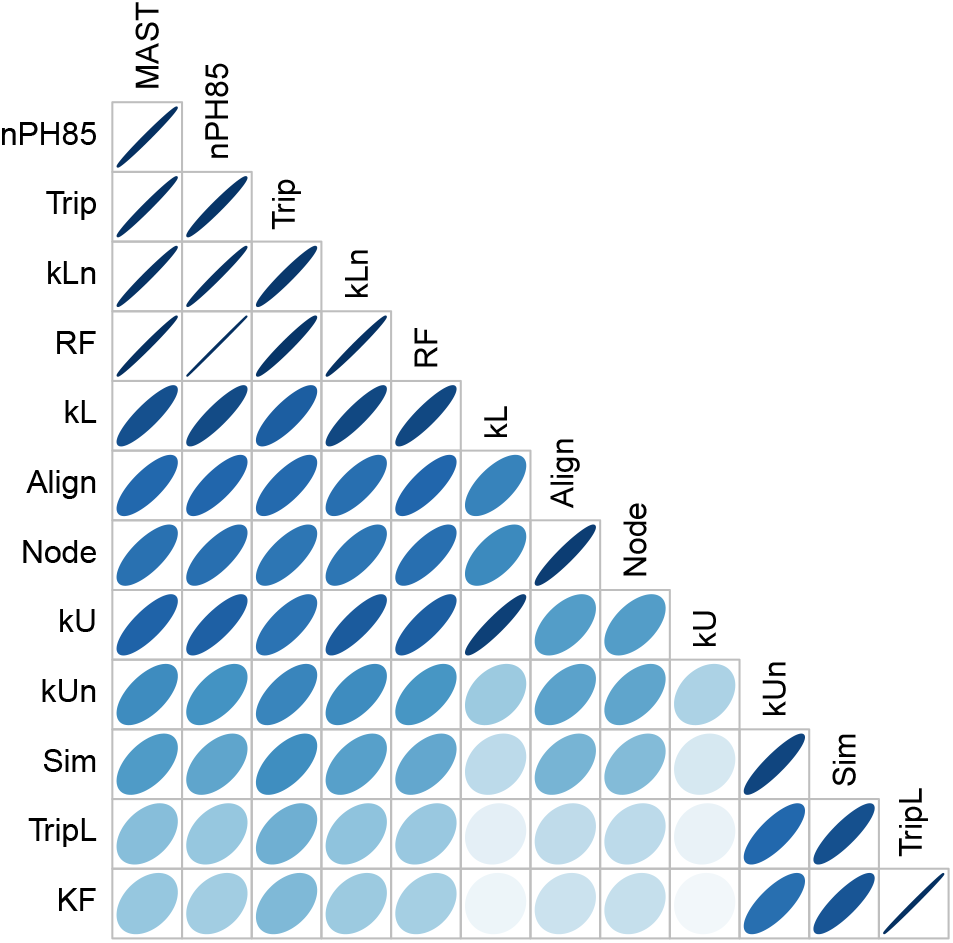
Summary of correlation matrix of tree distance measures. Spearman’s rank correlations were calculated for 10,000 trees simulated under varying model parameter settings, wider and blue-darker is the bubble, higher is the correlation.

We used the mutual information (MI) to quantify the information content of each metric with respect to the three model parameters. Based on our preliminary results with the edge case scenarios, we calculated a second set of MI values where the parameter space was constrained to *P* > 0.8 for *M* and Λ; and by *M* < 10 ^−4^/lineage/Ma for Λ and *P* (Fig. 4). Overall, Sim was the most informative metric for Λ, while kU and kL were the most informative for P. Several metrics obtained similar levels of MI for M, including kL, RF and nPH85.

**Figure 4.**
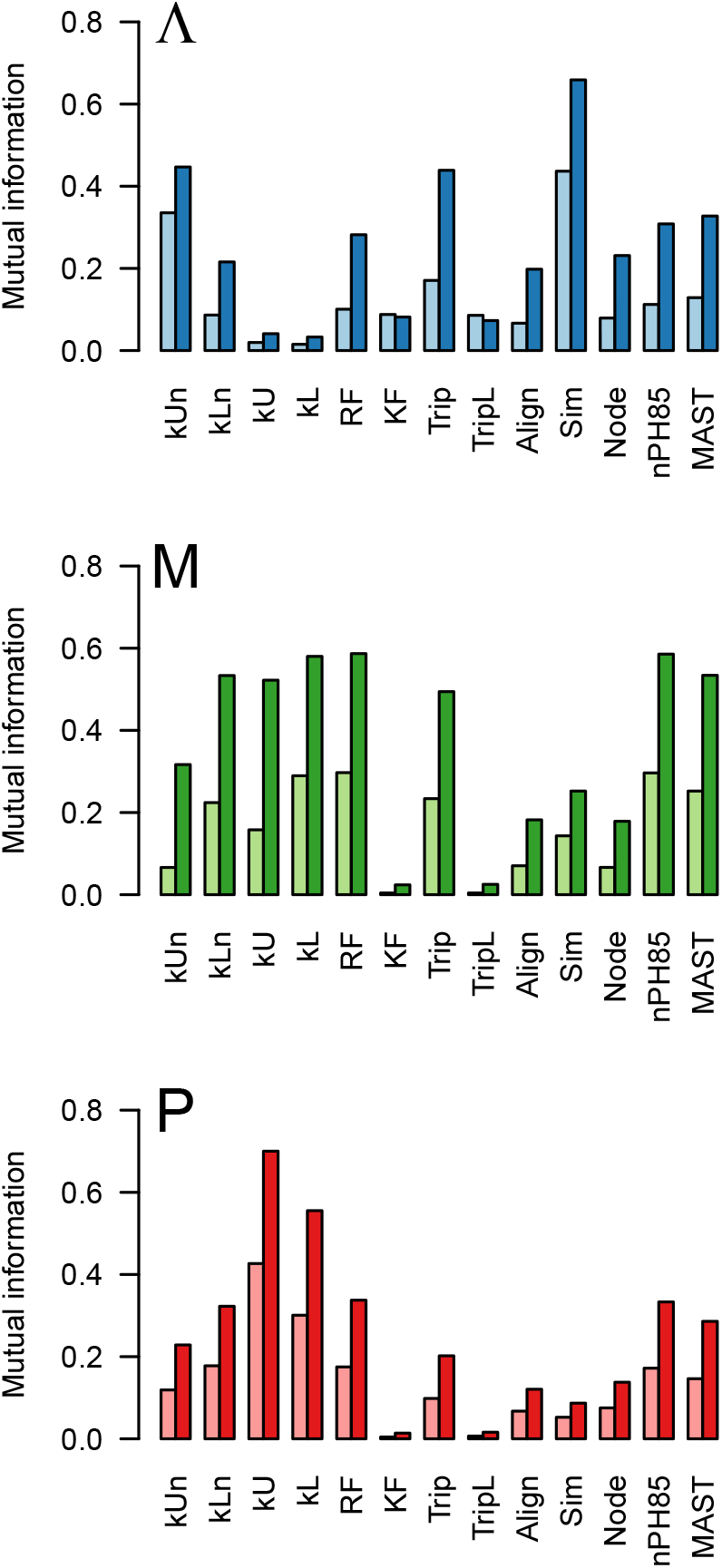
Barplots summarizing the mutual information of distance measures on the cophylogeny model parameters. The mutual information I was calculated by discretizing each distance *d* and model parameter *θ* into 10 bins respectively, for a total of 100 bins in the joint distribution *p*(*d*, *θ*), and then computing the sum 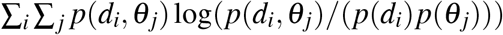. If *I* = 0, then *d* is independent of *θ*. Two values of *I* were computed for each distance. The left values were computed from the entire parameter space, whereas the right values were constrained as follows: (Λ) low coalescence, *M* < 10^-4^; (*M*) high cospeciation *P* > 0.8, and; (*P*) low migration, *M* < 10 ^-4^.

To examine the response of kU to variation in *P* and *M* more closely, we generated contour plots for this metric and the popular RF distance for comparison (Fig. 5). These plots clearly illustrate that the information content of either metric on P is dependent on the migration rate, and decays as M becomes too high. We note that unlike Figure 1, where the coalescence rate was fixed, these contour plots mask extensive variation in Λ among simulations. Similarly, Figure 6 illustrates the response of the metrics Sim and kUn to variation in Λ and *M*. Again the information content of either metric on Λ decays when *M* becomes too high; this effect is more conspicuous for Sim.

**Figure 5:**
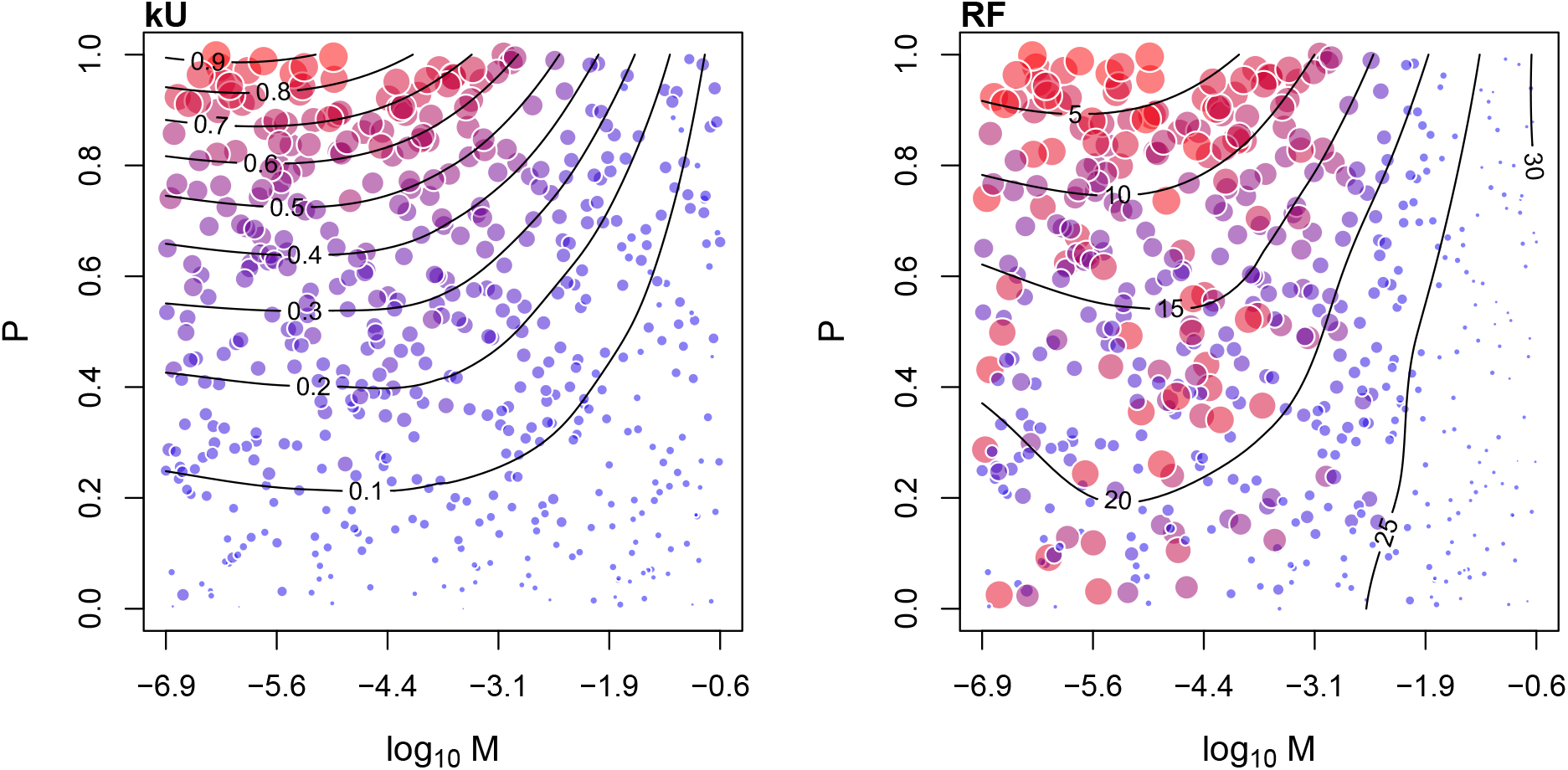
Contour plots summarizing the response of the unlabeled kernel (kU) and Robinson-Foulds distance (RF) to variation in cospeciation probability (*P*) and migration rate (*M*). Each point represents the average of 100 replicate simulations for a given parameterization of the cophylogeny model. The area and colouring of points is proportional to the distance metric.

**Figure 6:**
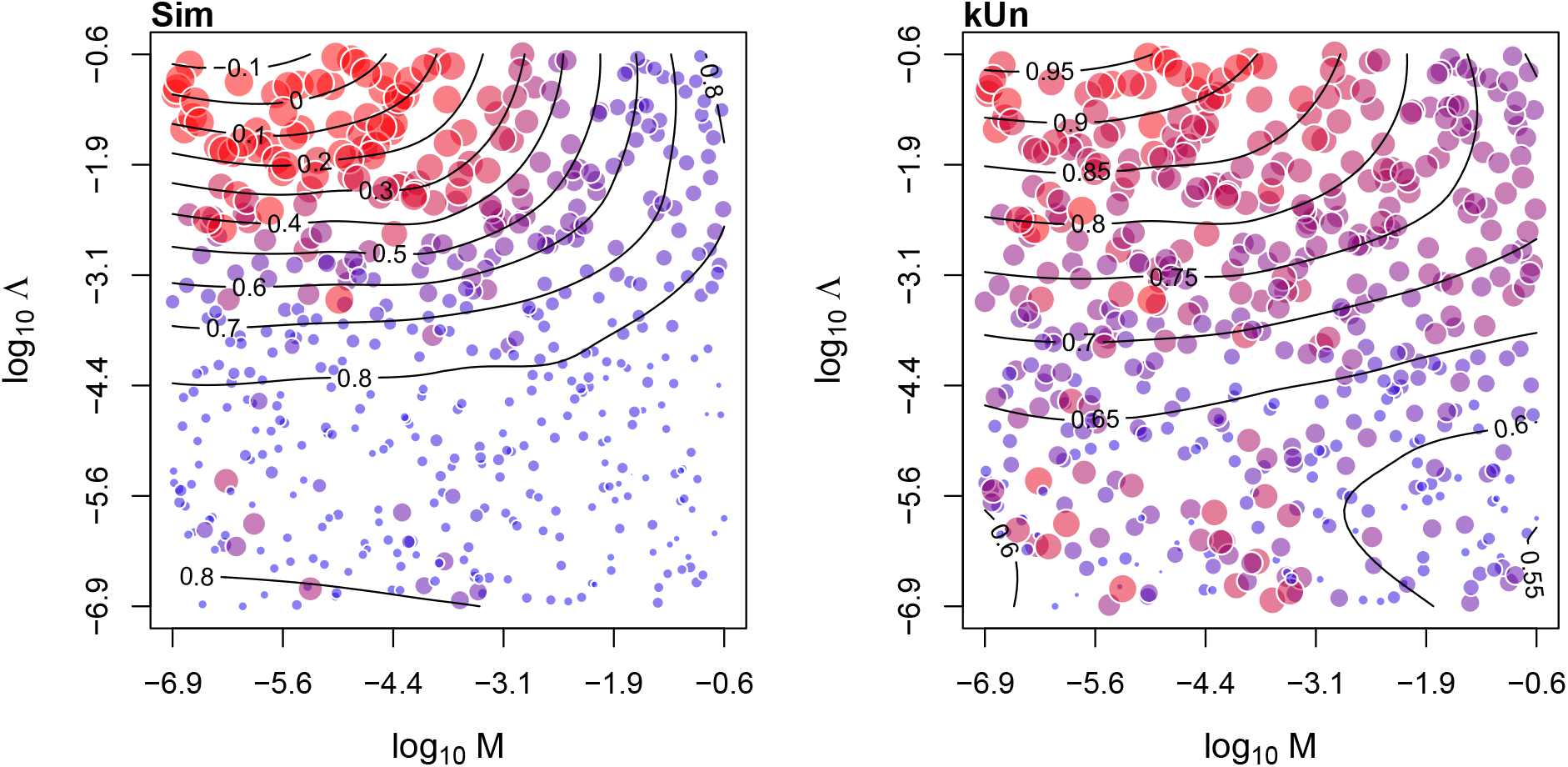
Contour plots summarizing the response of the Sim distance and normalized unlabeled kernel (kUn) to variation in coalescence (Λ) and migration (M) rates. Each point represents the average of 100 replicate simulations for a given parameterization of the cophylogeny model. The area and colouring of points is proportional to the distance metric.

### Application to real data sets

Our simulation experiments reveal that the different distance measures respond differently to variation in coalescence, migration and cospeciation rates. Furthermore, none of the distance measures is independently capable of conveying substantial information about all three cospeciation parameters. Although the simulated data provide a ‘ground truth’ to these parameters, the underlying model relies on unrealistic assumptions (see Discussion section) that limit the biological realism of these data. To assess the response of these distance measures to phylogenies reconstructed from actual data, we collected published trees or sequences for matched sets of host and pathogen species from the literature. We searched the literature for studies of host-pathogen cospeciation where the system was qualitatively described as having high or low levels of phylogenetic concordance due to cospeciation (the ‘General’ collection, Table 2). We used these descriptions to partition the ‘General’ collection into two categories.

This transition from simulated to actual data highlighted significant obstacles in the use of distance measures to cophylogeny studies. First, the metrics often require the trees to be the same size, *i.e.*, to have equal numbers of tips (Table 1). Distance metrics that utilize labels, such as the RF distance, also require that the trees have the same labels, *e.g.*, that the trees are alternative models for relating the same taxa. When simulating the data sets, it was trivial to generate pathogen trees that matched the labels of the host tree by initializing a single pathogen lineage in each host species. The biological reality of host-pathogen associations is frequently more complex, however. A pathogen species may be found in more than one host species, and a host species may be associated with multiple pathogen species. These cases may be accommodated by grafting additional branches with zero lengths to tips with multiple associations, to enforce a one-to-one map between the host and pathogen phylogenies (Geoghegan et al. 2017). Similarly, we grafted zero-length branches to equalize the numbers and labels of tips in each pair of trees in the ‘General’ data collection, and then calculated tree distance measures for each pair. When examining each distance measure individually, we did not observe any clear separation between high and low codivergence tree sets in the ‘General’ collection (Supplementary Material Fig. S1). This is consistent with findings from our simulation analysis that the distance measures vary substantially in their response to different cospeciation parameters. We next used a principal components analysis to examine the joint distribution of the general collection as a biplot (Fig. 7). This plot illustrates a rough separation between cases of high and low codivergence, with the exception of one low outlier (31-32) and a cluster of high outliers (1-2, 9-10, 21-22). Since the tree pair 31-32 has the fewest tips of any case (*n* = 5 for both hosts and parasites, Table 2), its placement in the biplot may simply be due to sampling variation. The low loadings on TripL and KF in the biplot are consistent with their low mutual information with respect to any cospeciation parameters in our simulations (Fig. 4). Similarly, the variable loadings on the first component of the biplot — combined with our simulation results — suggest that the characterization of phylogenetic concordance in these studies is strongly influenced by host switching (migration) events.

**Figure 7:**
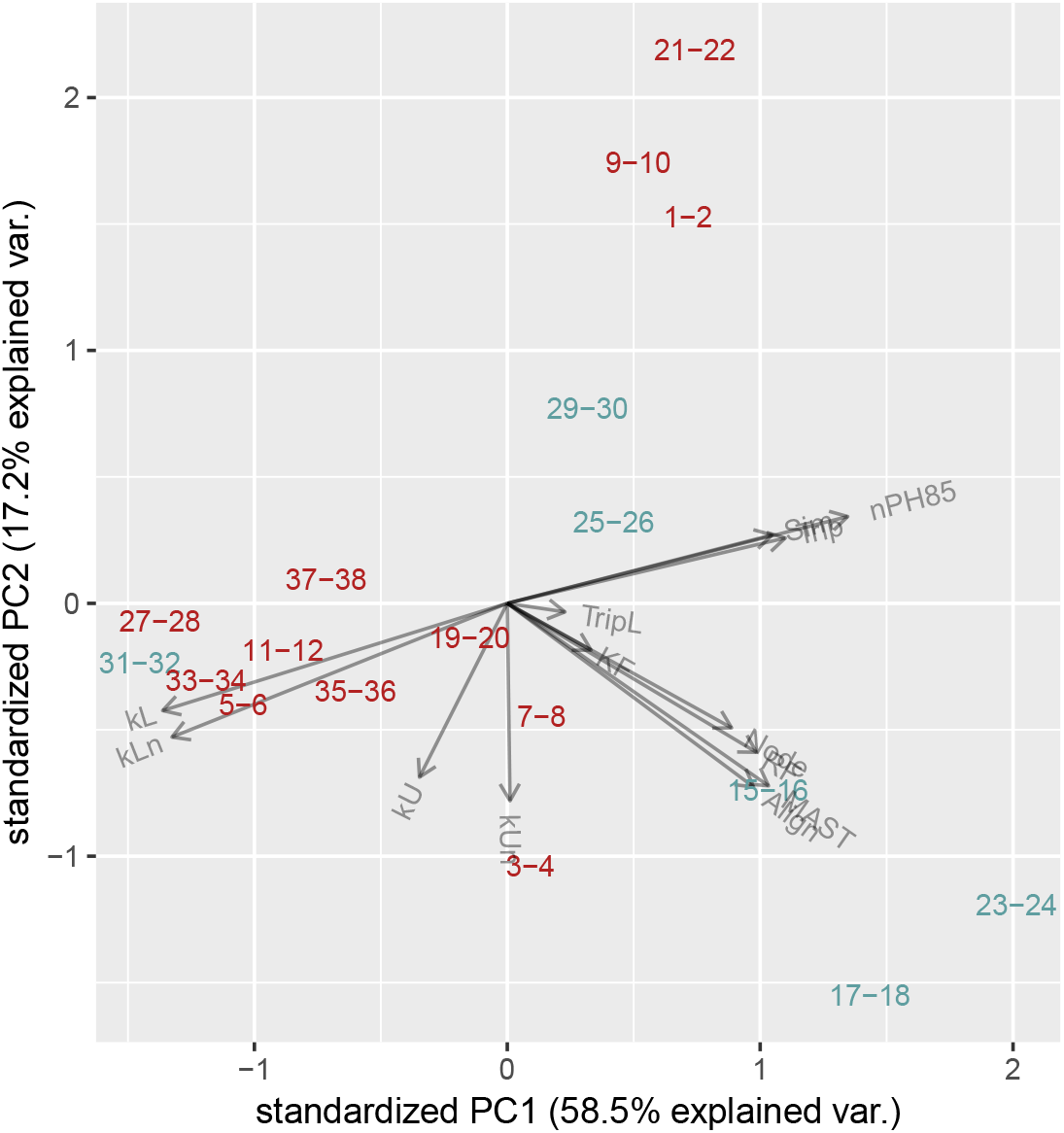
Biplot of a principal components analysis on distance measures for the general dataset. Each label represents a pair of host and pathogen trees that were characterized in the respective sources as cases of high (red) or low (blue) codivergence (see also Table 2). The gray vectors represent the variable loadings for the respective distance measures.

One of the unique features of the kernel methods compared to the other distance measures in this study is that they can be applied to unlabeled trees. This enables us to not only compute a distance between a pair of host and pathogen trees, but we can also compute distances between a host tree and pathogen trees from other pairings. In other words, it is not possible to compute the RF distance between the trees relating crinoids (sea lilies, Lanterbecq et al. (2010), from couple 21-22) and the gut bacteria of termites (Noda et al. (2007), from couple 37-38) even though these trees are the same size. We can therefore embed all the trees into a common feature space defined by a given kernel (Supplementary Material Fig. S2). We exploited this characteristic to test whether pairs of trees in the general collection were significantly closer together in this feature space than expected by chance with a randomization test. We drew 18 random pairings of host and pathogen trees from the ‘General’ collection, calculated the mean unlabeled kernel score (kU), and repeated this procedure to obtain 1,000 replicate means to approximate a null distribution. The mean kernel score for the actual tree pairs (*E*(*kU*) = 0.90) was located in the 99.9 percentile of this distribution, indicating that the actual pairs were significantly closer in the kU-defined space than expected by chance (*p* = 2.0 × 10^-4^).

Finally, we examined a second collection, the ‘Viral’, of host and pathogen trees corresponding to 19 different virus families from a previous study (Geoghegan et al. 2017). Since the host trees were derived from a common time tree, we generalized the host tip labels to the family taxonomic level, making it feasible to compare non-associated trees with both unlabeled and labeled kernels. In addition, branch lengths in the host trees were scaled in time to millions of years, whereas the pathogen trees were scaled to evolutionary time (expected numbers of nucleotide substitutions). This difference made it necessary to renormalize branch lengths in both host and pathogen trees (dividing by mean branch length) for kernel-mediated comparison (kUn and kLn). Supplementary Material Figure S3 comprises two PCA plots from the analysis of the similarity matrices using the unlabeled (kUn) and labeled (kLn) kernel functions, respectively. Again, we ran randomization tests for this collection using either kUn and kLn. When we ignored labels in comparing tree shapes, the mean kernel score for the actual tree pairs was *E*(kUn) = 0.78 and located in the 33.1 percentile (*p* = 0.67) of a randomized null distribution, indicating that the actual pairs were not significantly closer in the kUn-defined space than expected by chance. We obtained substantially different results with a labeled kernel: the mean score (E(kLn) = 0.19) was located at the 99.9 percentile of the randomized distribution (*p* = 7.0 × 10^-4^), indicating that the actual pairs were significantly closer in this feature space than expected by chance. Geoghegan et al. (2017) previously reported that the phylogenies of DNA viruses and their hosts tended to be more concordant than RNA viruses, which was attributed to their relatively higher rates of cospeciation and lower rates of migration. Here we observed the same trend for families of DNA viruses, especially Hepadnaviridae, Poxviridae and Papillomaviridae. However, we also observed significant clustering for the RNA virus families Orthomyxoviridae and Potyviridae. In the latter case, clustering was most likely driven by the unique distribution of these viruses in plant host species. Using non-parametric Wilcoxon tests, we found no significant difference in kUn distances separating DNA or RNA virus trees from their respective host trees (*p* = 0.17), but significantly greater labeled (kLn) distances for RNA viruses (*p* = 0.02).

## Discussion

There is a deep literature on developing distance measures for the comparison of phylogenetic trees in order to quantify biological processes such as speciation (Mooers 1997; Kuhner and Yamato 2014). Multiple quantitative frameworks for the comparison of phylogenetic trees have also been developed for the study of cophylogeny, to determine whether the two sets of organisms share a coevolutionary history (*e.g.*, Huelsenbeck et al. 2000; Doyon et al. 2011). We therefore anticipated extensive applications of tree distances in the literature for analyzing cophylogeny or coevolution. However, our survey on papers citing the tree distance metrics found only four studies that have made use of metrics for coevolutionary studies (0.001% of all studies reviewed, Table 1), which is a surprising outcome given the similar objectives of the respective fields. Instead, the comparison of trees in coevolutionary studies had frequently relied on other methods where a tanglegram is either assessed qualitatively by the investigator, or analyzed with a reconciliation method, which is computationally complex and requires a subjective assignment of cost functions to the respective coevolutionary events (*e.g.*, cospeciation, host switching and extinction). The objective of our study was to assess the potential utility of distance metrics for cophylogenetic studies.

In this study, we did not attempt to evaluate a comprehensive set of all available distance metrics. Instead we have focused on a subset of metrics from a recent review by Kuhner and Yamato (Kuhner and Yamato 2014), augmented with a small number of additional metrics including the kernel metrics. In addition to the metrics in our study, there is a large number of distance metrics that can be constructed from summary statistics such as Sackin’s index (Blum and François 2005). A summary statistic reduces a tree down to a single number that quantifies a biologically significant aspect of tree shape such as asymmetry (Mooers 1997). Thus, we can obtain a distance from a summary statistic by taking its difference between the host and pathogen trees. However these summary statistics usually do not incorporate tip labels, placing greater emphasis on similarity in tree shapes, and they can be difficult to normalize for comparing pairs of trees with different sizes (Pompei et al. 2012). In addition, there are several spectral methods that can be applied to trees by interpreting these objects as graphs (Hendy and Penny 1993) and the further development of tree distances continues to be an active area of research (Kendall and Colijn 2016; Colijn and Plazzotta 2017). It is therefore not feasible to evaluate all possible distances and for our purposes, we have only evaluated a representative subset of distance measures, including commonly used measures such as the RF distance.

Simulation experiments are an essential step to evaluate the response of a metric to variation in the data because the underlying parameters are known without ambiguity. However the inherent assumptions of the simulation model may limit our ability to extrapolate from that analysis to real applications. In this study, we have taken the unusual approach of simulating the pathogen trees backwards in time along a fixed host tree. Our motivation for this approach is that it is more efficient to start from ‘sampled’ lineages and converge back in time to their common ancestors, then to simulate forward from a single ancestor and discard cases that are not compatible with the expected endpoints. For example, a forward-time simulation may simply go extinct before any lineages can become sampled. This approach makes it difficult to incorporate unobserved extinction events, although inferring these events is already difficult due to the sensitivity of extinction rate estimates to model misspecification (Rabosky 2010). In addition, we made a simplifying assumption that a single pathogen lineage was sampled per host. Although it is straight-forward to model the sampling of multiple pathogen lineages in a host species within our reverse-time framework, we sought to minimize the complexity of the parameter space to evaluate in our simulation experiments. Similarly, we assumed complete sampling of all extant pathogen lineages.

Our simulation analyses of tree distance metrics demonstrated that some metrics were more informative than others with respect to specific cospeciation parameters. For example, the Sim metric (Hein et al. 2004) was the most responsive to variation in coalescence rates, and the un labeled kernel (kU) to variation in cospeciation probabilities. The metrics evaluated in this study were frequently correlated with each other, but the correlations are seldom so extreme that the metrics were essentially redundant, *e.g.*, the RF and nPH85 distances. We also determined that distance metrics were more informative about the model parameters when the underlying parameter values were not so extreme that the host tree has essentially no influence on the shape of the pathogen tree; i.e., when the migration (host switching) rate was not too high, or when the cospeciation probability was substantially less than one and the pathogen coalescence rate was near zero (Fig. 1). These scenarios would make it difficult to meaningfully quantify cophylogeny by any method. If the host switching rate is exceedingly high, then the pathogen species are ‘cosmopolitan’ and freely utilize whichever host species they encounter, which would negate any influence of cophylogenetic effects on the pathogen phylogeny. In the second scenario, the pathogen speciation rate is so low that pathogen lineages speciate on a much longer time scale than their hosts, making the distribution of coalescent events independent of the host phylogeny. This scenario may arise when pathogen gene flow is unrestricted among host species (Johnson et al. 2003).

Next, we applied these metrics to two collections of phylogenies that were reconstructed from actual biological data. In the ‘General’ collection of coevolutionary studies across all taxonomic groups, we retrieved a total of 18 studies — including parasitic and symbiotic associations — where authors described the trees as having a high or low degree of concordance. Only 6 of these studies reported low concordance. These assignments were largely based on a subjective qualitative assessment of phylogenetic concordance, and there are no quantitative criteria that have been applied generally across taxa. Given the broad diversity of taxonomic groups being studied, it is unlikely that any one of the coevolutionary processes is consistently determining either outcome. It is also not feasible to determine with complete certainty how each process contributed to the varying levels of concordance across these empirical studies. Nevertheless, the projection of these trees into a parameter space defined by the 13 distance metrics revealed some clustering of studies reporting low concordance. This suggests that different investigators are describing concordance in a consistent way across biological systems, and that these subjective assessments can be at least partly quantified using distance metrics. In Figure 7, we observed a small cluster of studies, where the authors reported high concordance between host and pathogen trees (1-2, 9-10 and 21-22), that did not associate with the main grouping of ‘high concordance’ studies. We found that these studies may have placed greater emphasis on a relative lack of crossed association edges in the tanglegrams that implies a low rate of host switching. However, the topologies of the trees in the respective studies are markedly dissimilar in shape, such that we believe these studies should instead be classified as examples of low concordance. Our analysis suggests that this discordance in these cases may be driven by a lower probability of cospeciation, based on our assessment of the tanglegrams in the respective publications, the alignment of this cluster with the kU vector, and our simulation results.

Reconciliation methods implicitly assume that the pathogen phylogeny is the outcome of a stochastic process that has unfolded along the host phylogeny, shaped by events such as cospeciation or migration that have occurred at different rates. The distance-based approach that we have evaluated in this paper is analogous to fitting a non-parametric model to the shape of the pathogen phylogeny, conditional on the host phylogeny — none of these processes is explicitly modeled by any of the distances evaluated in this study. Although many methods employ maximum parsimony to infer these events, the problem of reconciliation lends itself to probabilistic inference through maximum likelihood (Huelsenbeck et al. 1997) and Bayesian (Huelsenbeck et al. 2000) frameworks, which have already been developed for restricted scenarios, *e.g.*, no speciation within hosts (Paterson and Banks 2001). The ideal Bayesian approach would be to jointly sample the host and pathogen phylogenies and reconstructions of coevolutionary events given the sequence and associational data — however, the enormous model space this would entail would likely limit this approach to small data sets. There is growing interest across disciplines in using simulation-based methods, *e.g.*, approximate Bayesian computation (ABC), to estimate parameters instead of directly calculating model likelihoods (Tavaré et al. 1997). The basic premise of ABC is that fitting can proceed by adjusting the parameters of the model until it yields simulations that resemble the observed data. Although ABC is intuitively appealing and relatively straight-forward to implement, it is challenging to find similarity measures for comparing simulated and observed trees that are efficient to compute and sufficiently informative to estimate the parameters. Baudet et al. (2014) recently used an ABC approach to cophylogeny using forward-time simulation of pathogen trees on a fixed host phylogeny, and employed a single distance metric based on the number of tip labels shared between the largest isomorphic subtrees. Their results indicated a general lack of parameter identifiability, such that a given pair of trees can be explained equally well by a broad range of event combinations. In another recent paper, Alcala et al. (2017) applied multiple network statistics (*e.g*., degree size) to simulated tanglegrams to estimate host switching and cospeciation rates using a rejection ABC method. We anticipate that the analysis of distance metrics presented here will provide an important foundation for the further development of ABC-based methods as a promising approach to the study of cophylogeny.

## Acknowledgements

This work was supported in part by the Government of Canada, through Genome Canada and the Ontario Genomics Institute (OGI-131), and by grants from the Canadian Institutes of Health Research (PJT-155990, PJT-156178) and the Natural Sciences and Engineering Research Council of Canada (RGPIN-2018-05516). We also thank Eric Wong and Vidhu Joshi, for their assistance in searching the literature for cophylogeny studies and collecting published data sets, Faisal Abu-Sardanah and David W. Dick for technical support.

## Supplementary Material Figures

**Figure S1:**
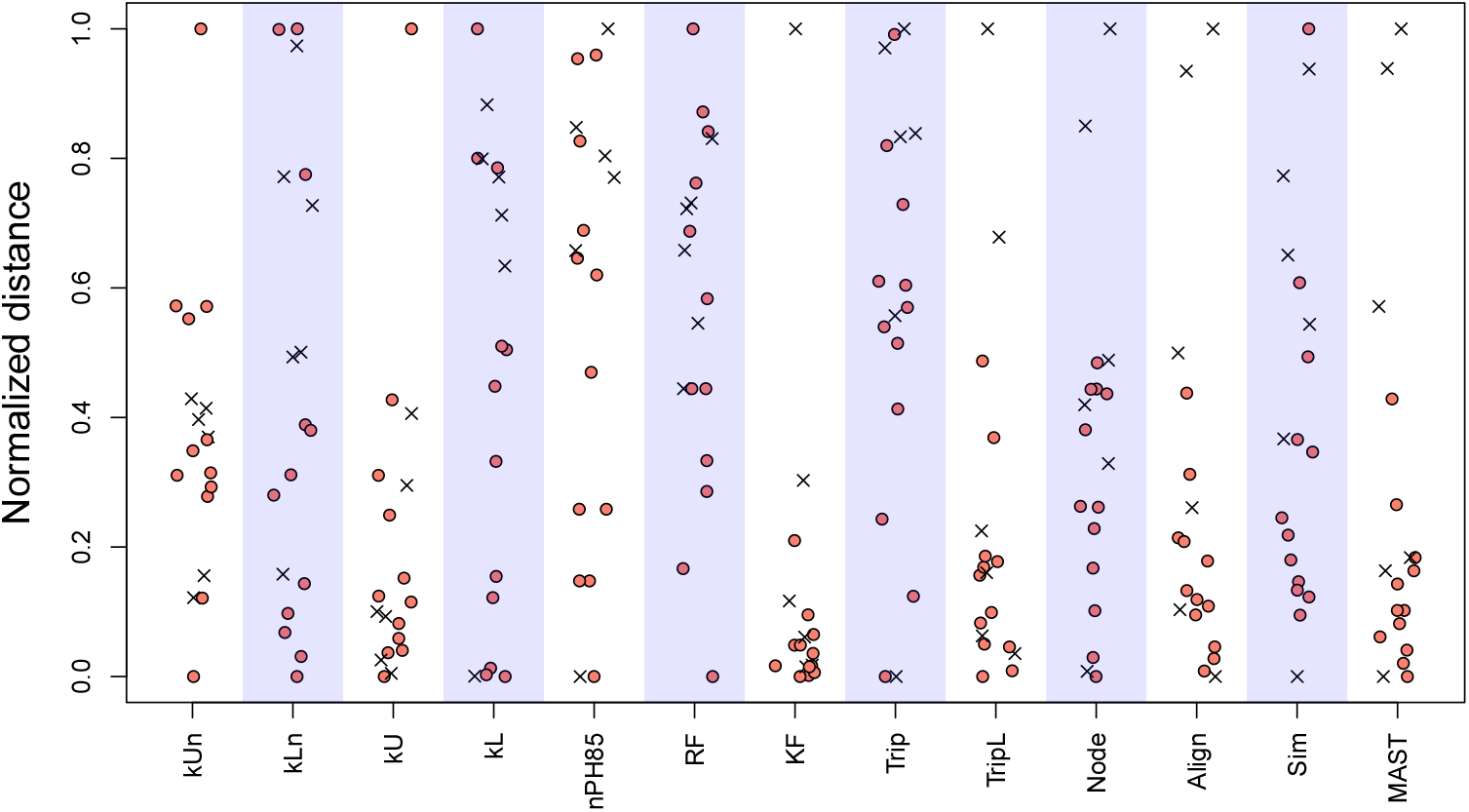
Tree distance measures for the general collection of host-pathogen trees. High- (circles) and low-codivergence (crosses) assignments were drawn from the publications associated with each pair (Table 2). We grafted zero-length branches to tips with multiple relations, *e.g.*, a pathogen species found in two host species, to enforce a one-to-one relation between tips in the host and pathogen trees. To facilitate comparisons across distance measures, each measure x was normalized to range from 0 to 1 by applying the transformation (*x* — min(*x*))/ (max(*x*) — min(*x*)).

**Figure S2:**
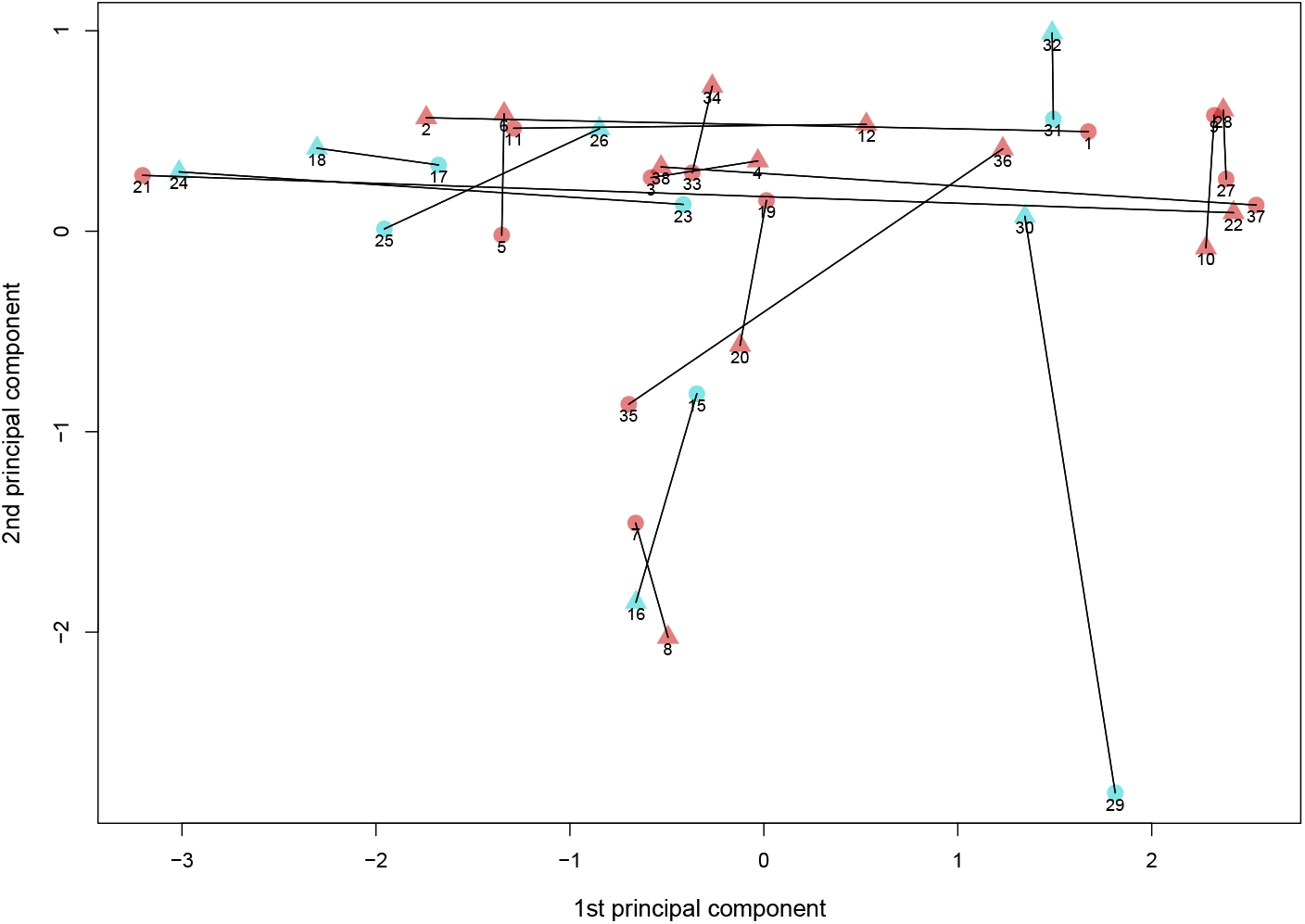
PCA on ‘General’ collection unlabeled kernel (kU) scores showing host and pathogen pairs coming from high co-divergence (in red) and low-codivergence (in cyan) studies as summarized in Table 2. Every single point represents host or pathogen tree’ eigenvalue, with the one belonging to the same pair united by a straight line.

**Figure S3:**
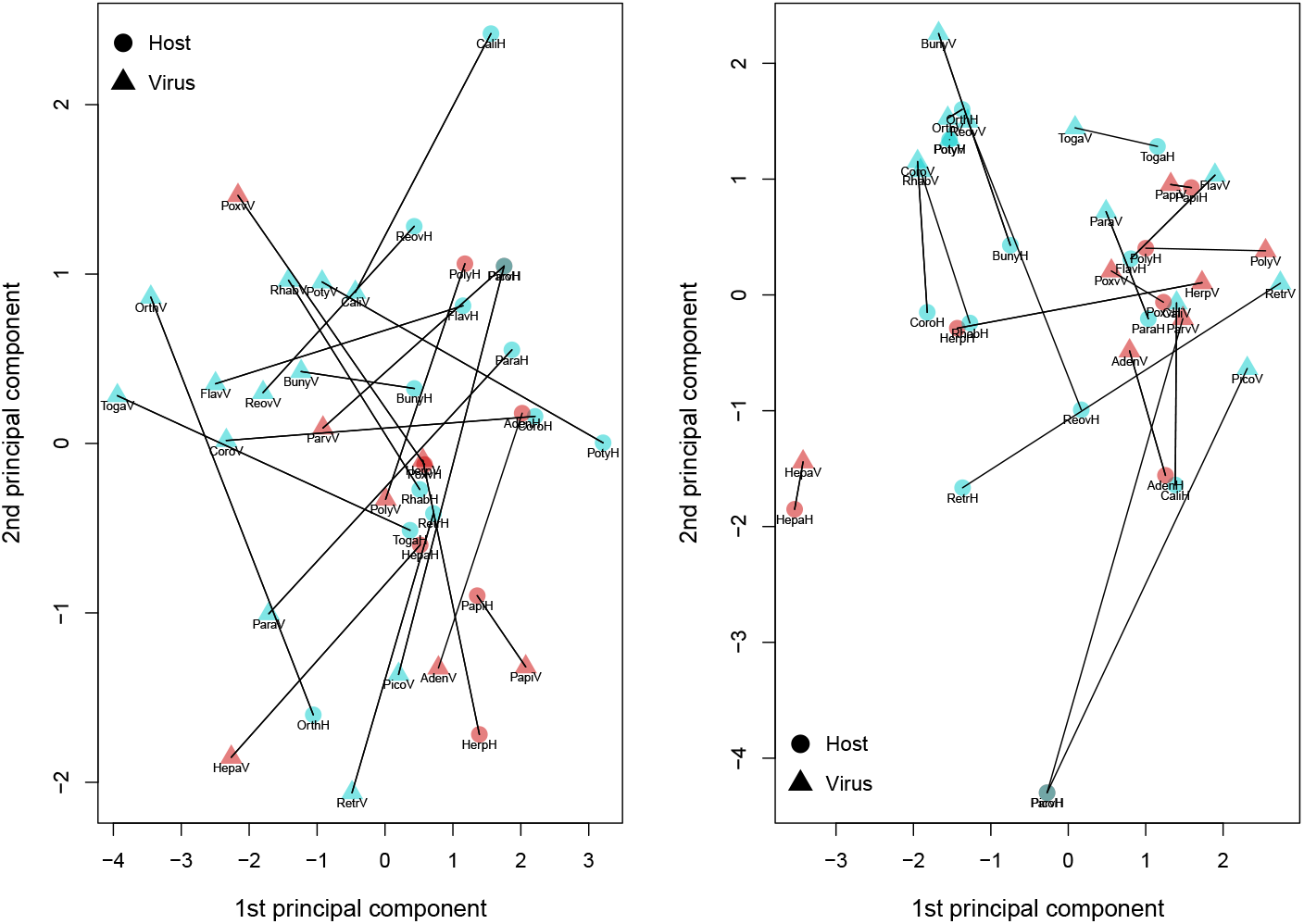
PCA for 19 virus families, a: kUn; b: kLn. In red pairs including DNA-viruses, in cyan pairs icluding RNA-viruses. Aden: Adenoviridae, Buny: Bunyaviridae, Cali: Caliciviridae, Coro: Coronaviridae, Flav: Flaviviridae, Hepa: Hepadnaviridae, Herp: Herpesviridae, Orth: Orthomyxoviridae, Papi: Papillomaviridae, Para: Paramyxoviridae, Parv: Parvoviridae, Pico: Picornaviridae, Poly: Polyomaviridae, Poty: Potyviridae, Poxv: Poxviridae, Reov: Reoviridae, Retr: Retroviridae, Rhab: Rhabdoviridae, Toga: Togaviridae, H: Host tree, V: Virus tree, host and virus’ trees have same symbol.

## Supplementary Material Tables

**Table S1:** Parameter settings used for simulation experiments corresponding to three edge cases. Each edge case varied one of the model parameters over the range 0 to 1 inclusive. In addition to these endpoints, we selected 10 to 12 arbitrarily chosen intermediate values. For each parameter combination, we simulated 100 replicate pathogen trees.

| Edge case | $\Lambda$ | $M$ | $P$ |
| --- | --- | --- | --- |
| Coalescence only | $10^{-6}, 10^{-5}, 10^{-4},$<br>0.001, 0.01, 0.025,<br>0.04, 0.07, 0.13, 0.25,<br>0.5, 0.75, 0.9, 1 | 0 | 0 |
| Migration only | 1 | 0, $10^{-5}, 10^{-4},$<br>$5.5 \times 10^{-4}, 10^{-3},$<br>$2.125 \times 10^{-3}, 3.25^{-3},$<br>$4.375 \times 10^{-3},$<br>$5.5 \times 10^{-3}, 0.01,$<br>0.25, 0.5, 0.75, 1 | 1 |
| Cospeciation only | $10^{-6}$ | 0 | $10^{-6}, 10^{-5}, 10^{-4},$<br>0.001, 0.01, 0.1, 0.25,<br>0.5, 0.75, 0.9, 0.99, 1 |

